# Molecular dissection of carbapenem-resistant *Acinetobacter baumannii* circulating in Indian hospitals using Whole Genome Sequencing

**DOI:** 10.1101/2021.07.30.454432

**Authors:** Steffimol Rose, Varun Shamanna, Anthony Underwood, Geetha Nagaraj, Akshatha Prasanna, Vandana Govindan, Sravani Dharmavaram, M R Shincy, Silvia Argimón, Monica Abrudan, David M Aanensen, K. L. Ravikumar

## Abstract

**Objectives:** Carbapenem-resistant *Acinetobacter baumannii* (CRAB) has acquired worldwide recognition as a serious nosocomial infection. It poses a concern to hospitalized patients because of the limited therapeutic options available. Thus, we investigated the molecular epidemiology and antibiotic resistance profiles of *A. baumannii* isolates in India.

**Materials and Methods:** We characterized 306 retrospective *A. baumannii* clinical isolates collected from 18 centers across 10 states and 1 Union Territory of India between 2015 and 2019. Molecular epidemiology, and carbapenem resistance were studied by Whole Genome Sequencing.

**Results:** A total of 105 different Sequence Types (STs) were identified including 48 reported STs and 57 Novel STs. 99 isolates were classified into Clonal Complex 451 (CC451) among which ST848 and ST1956 were the common STs. Carbapenemase resistance was confirmed in all the isolates with the presence of intrinsic *bla*_OXA-51-like_ genes, and the acquired *bla*_OXA-23_ and *bla*_NDM-1_ genes.

**Conclusion:** Most of the isolates were grouped under clonal complex 451. ST1053 caused an outbreak in Northern India during 2018 and 2019. Novel MLST alleles and STs were also detected, underlining an evolutionary divergence in India. The carbapenem-resistance was dominated by OXA-type carbapenemases and further surveillance of these carbapenem-resistant *A. baumannii* and antimicrobial stewardship should be strengthened.

## Introduction

*Acinetobacter baumannii* has emerged as the most common cause of hospital-acquired illnesses**[1]**. These non-fermentative, gram-negative coccobacilli are a frequent cause of hospital outbreaks, particularly in patients in intensive care units **[2]**. Infections due to *A. baumannii* includes ventilator-related pneumonia, sepsis, urinary tract infections, skin, and soft tissue disorders resulting in greater mortality and morbidity due to the scarcity of therapeutic options, intrinsic resistance to toxins and antibiotics, transformability, metabolic versatility, and genome plasticity **[3, 4]**. Although this bacteria has demonstrated its ability to survive in a variety of environments, the natural reservoir is yet to be discovered **[5]**. In recent years, hospitals in India have observed an increase in infections caused by gram negative bacteria including *A. baumannii* infections **[6]**.

It is vital to characterize the molecular epidemiology of *A. baumannii* isolates engaged in nosocomial infections in order to prevent their spread in the hospital environments. Multi Locus Sequence Typing (MLST) based on sequence analysis of loci from housekeeping genes is a frequently used method for pathogen typing**[7]**. MLST allows for inter-laboratory comparison, making it a useful tool for epidemiological research around the world **[8]**. There is a scarcity of published information about Sequence Types (ST) of Indian isolates. This emphasises the importance of routine sequence typing of *A. baumannii* strains in India, as well as the submission of STs to databases to track their epidemiology.

*A. baumannii* has a remarkable ability to acquire resistance to multiple antibiotics, especially carbapenems and it has been reported worldwide **[9, 10]**. Carbapenems are one of the most commonly used antibiotics to treat *Acinetobacter* infections and are in fact one of the last resort treatments. However, according to previous studies between 2013 and 2016, carbapenem resistance in India increased from 55% to 90.5% **[11]** making it a growing therapeutic concern. The production of carbapenemases (class D enzymes or oxacillinases and class B enzymes or metallo-β-lactamase), overexpression of efflux pumps, genetic changes of penicillin binding proteins (PBP), and loss of porin expression are all factors that contribute to increased carbapenem resistance **[12]**. Carbapenem resistance caused by OXA type enzymes include naturally occurring *bla*_*OXA–51-like*_ and three acquired genes: *bla*_*OXA–23*_, *bla*_*OXA-24*_, and *bla*_*OXA-58*_**[13]**.

Characterizing the *A*.*baumannii* isolates involved in nosocomial infections helps to monitor their spread in hospital settings and understand local epidemiology. In the study, Whole Genome Sequencing (WGS) was used to determine the genetic and epidemiological relatedness, and their prevailing resistance mechanisms of Indian hospital isolates. Investigating the clonal nature of these populations provides valuable data for infection management programs **[14]**.

## Materials and Methods

### Bacterial strains

Retrospective collections of non-repetitive Carbapenem-resistant *A*.*baumannii complex* isolates (n=341) were obtained from multiple hospitals in India during the years 2015-2019. The sample collections were from several backgrounds of infection and included isolates from different sources; sputum (n=101), tracheal aspirate (n=84), pus (n=77), blood (n=18), wound (n=15), urine (n=14), tissue (n=9), bronchoalveolar lavage (n=8), cerebrospinal fluid (n=6), catheter (n=4), swab and biopsy (n=2), fluid (n=1) **(Supplementary Table a)(Supplementary Fig a)**. Of the 341 *Acinetobacter* isolates included in this study, 90 were from Delhi, 111 from Karnataka (Kolar, n=70, Bangalore, n=23, and Mangalore, n=18), 52 from Pondicherry, 44 from Uttarakhand (Rishikesh), 14 from Uttar Pradesh (Lucknow), 10 from Bihar (Patna), 7 from Chennai, 6 from Rajasthan (Jaipur), 5 from Andhra Pradesh, and 2 from Maharashtra (Mumbai). The isolates were identified to the species level using Vitek2 (N281 Card, bioMérieux) and confirmed genotypically.

### Species Identification and Multilocus Sequence Typing

Genomic DNA was extracted from bacterial isolates with Qiagen QIAamp DNA Mini kit, in accordance with the manufacturer’s instructions. Double-stranded DNA libraries with 450 bp insert size were prepared and sequenced on the Illumina platform with 150 bp paired-end chemistry. Sequence reads generated in FastQ format were trimmed using Trimmomatic v.0.36 **[15]**, and assembled with SPAdes 3.15.0 **[16]**. The assemblies were run through the species identification tool BactInspector **[17]**. The isolates were sequence-typed with the MLST Oxford scheme **[18]** using the 7 house-keeping genes and assigned to an allele number. The allele numbers were combined to yield a specific ST utilizing the pubMLST database(https://pubmlst.org/abaumannii/). The goeBURST algorithm in PHYLOViZ **[19]** software was applied to assign STs to Clonal Complexes (CCs). Those sharing identical alleles at six of seven loci were interpreted as single locus variants (SLVs).

### Phylogenetic tree construction

SNP-based phylogeny was generated by mapping reads to a reference sequence using BWA mem; variants were called and filtered using bcftools, and a maximum likelihood phylogeny was produced using IQTree**[20**,**21**,**22]**. The tree was further visualized using Microreact**[23]**.

### Detection of carbapenemase genes

Resistome analysis to detect carbapenem resistance determinants for the assembled whole-genome sequences of the isolates was carried out using Ariba **[24]** tool and NCBI-AMR database using the protocols as detailed in www.protocols.io.

## Results

### Phenotypic characterization

The 341 *Acinetobacter* isolates were phenotypically identified as the *A. baumannii complex* by Vitek 2. 331/341 isolates passed the sequencing QC and were analyzed further. WGS distinguished the isolates into the following species, *A. baumannii (n=306), A. pitti (n=15), A. lactucae (n=8), A. radioresistans (n=1), and A. junii (n=1)*. 306 isolates belonging to *A. baumannii* were further analyzed for MLST.

### MLST analysis

Analysis of multilocus STs of the 306 *A. baumannii* isolates identified 49 different STs. The predominant ST was ST1053 (15%), followed by ST231 (5%), ST1956 (4%), and ST848 (4%). Other STs observed in the study are tabulated **(Table 1)(Supplementary Fig b)**. 57 Novel STs were detected in our study and submitted to the PUBMLST site for ST assignment. Single, double, and triple locus variants (SLV, DLV, TLV) of the 57 Novel STs were determined and are also tabulated in **Table 1**.

**Table 1:**
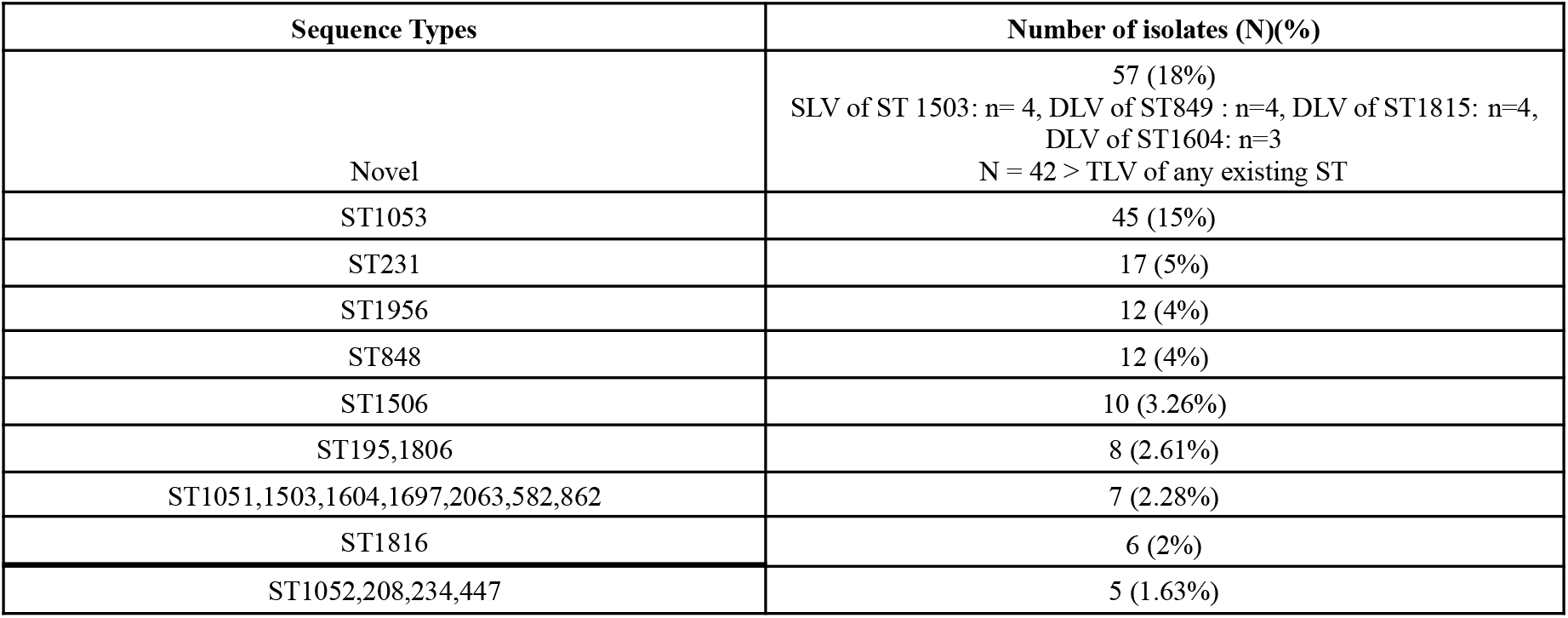
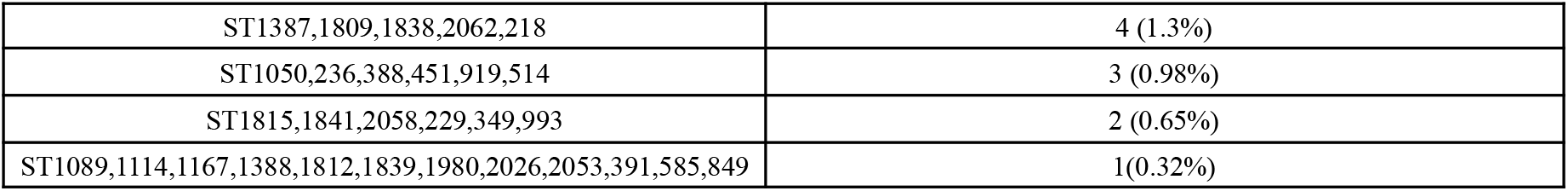
Summary of the sequence types observed in the study

### Clonal Complex analysis

The goeBURST analysis with PHYLOViZ software was performed to assign each ST to the respective clonal complex as represented in **Fig 1**. The goeBURST generated 4 major Clonal Complexes (CCs) (circled in Fig 1) with related STs. All the STs within this study grouped by CC are represented in **Fig 2 (Supplementary Table b)**. SLVs and DLVs of the STs in each of the CC have been labeled as 1 and 2 respectively in **Fig 1**. Seven STs, (ST234, ST236, ST514, ST1167, ST1089, ST1980, ST2053) were found as singletons and not related to any of the other STs in our dataset. SLVs and DLVs of all the STs identified in our study have been tabulated **(Supplementary Table c)**.

**Fig 1:**
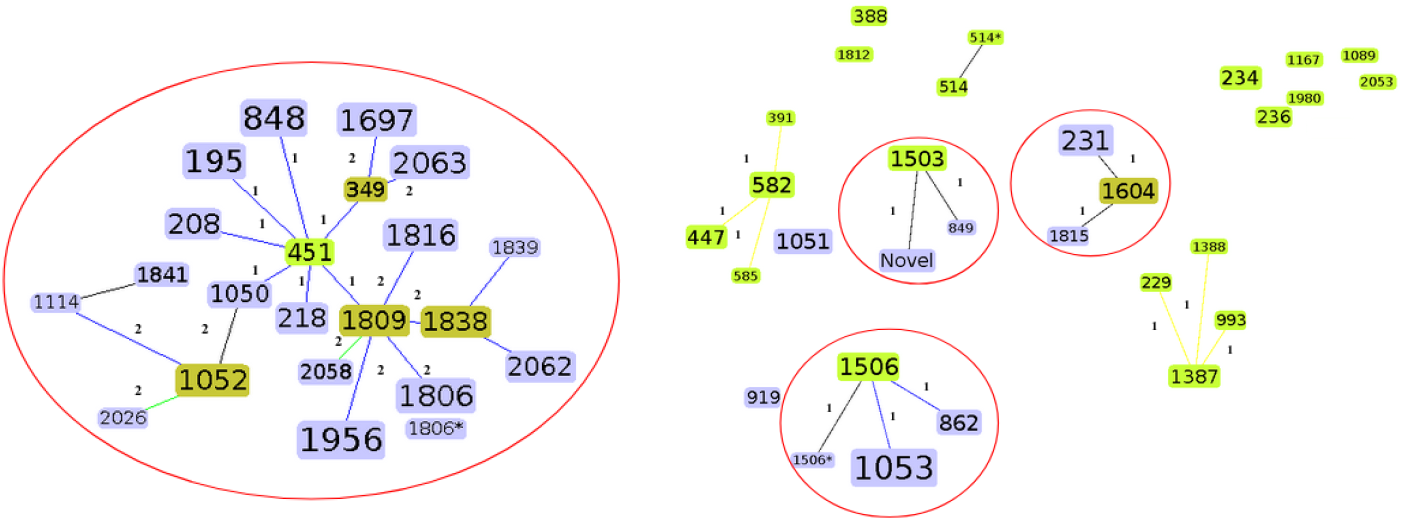
Clonal complex analysis by goeBURST. Each square signifies the sequence type. The size of each square represents the number of isolates, with larger sizes corresponding to the higher frequency of occurrence. Numbers 1 and 2 represent SLV and DLV.

**Fig 2:**
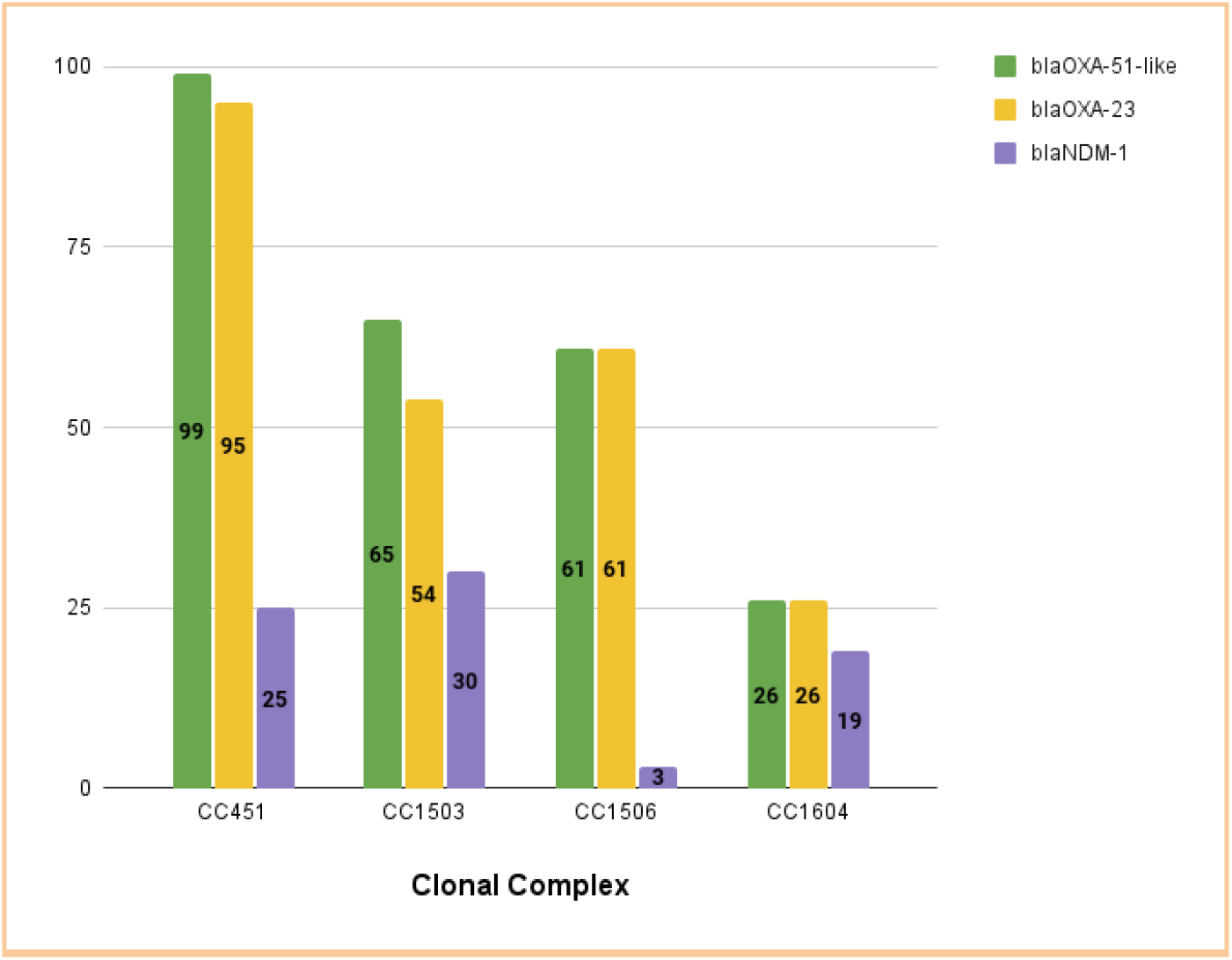
Figure describing STs within this study grouped by Clonal Complex and the Carbapenem resistance genes.

### Carbapenem Resistance

All 306 *A*.*baumannii* isolates carry the *bla*_OXA-51-like_ gene which is intrinsic to it. The acquired gene, *bla*_*OXA–23*_, belonging to the class D carbapenemases was observed in 92% (282/306) of the isolates. Interestingly, none of our isolates carried *bla*_*OXA-24*_, *and bla*_*OXA-58*_ genes. 29% of the isolates were found to be carrying the *bla*_*NDM-1*_ gene **(Table 2)**. Carbapenem resistance was observed in 97% (299/306) isolates which carried *bla*_*OXA-51-like*_, *bla*_*OXA–23*,_ and *bla*_*NDM-1*_ genes, while the remaining 3% isolates conferred resistance to beta-lactams.

**Table 2:**
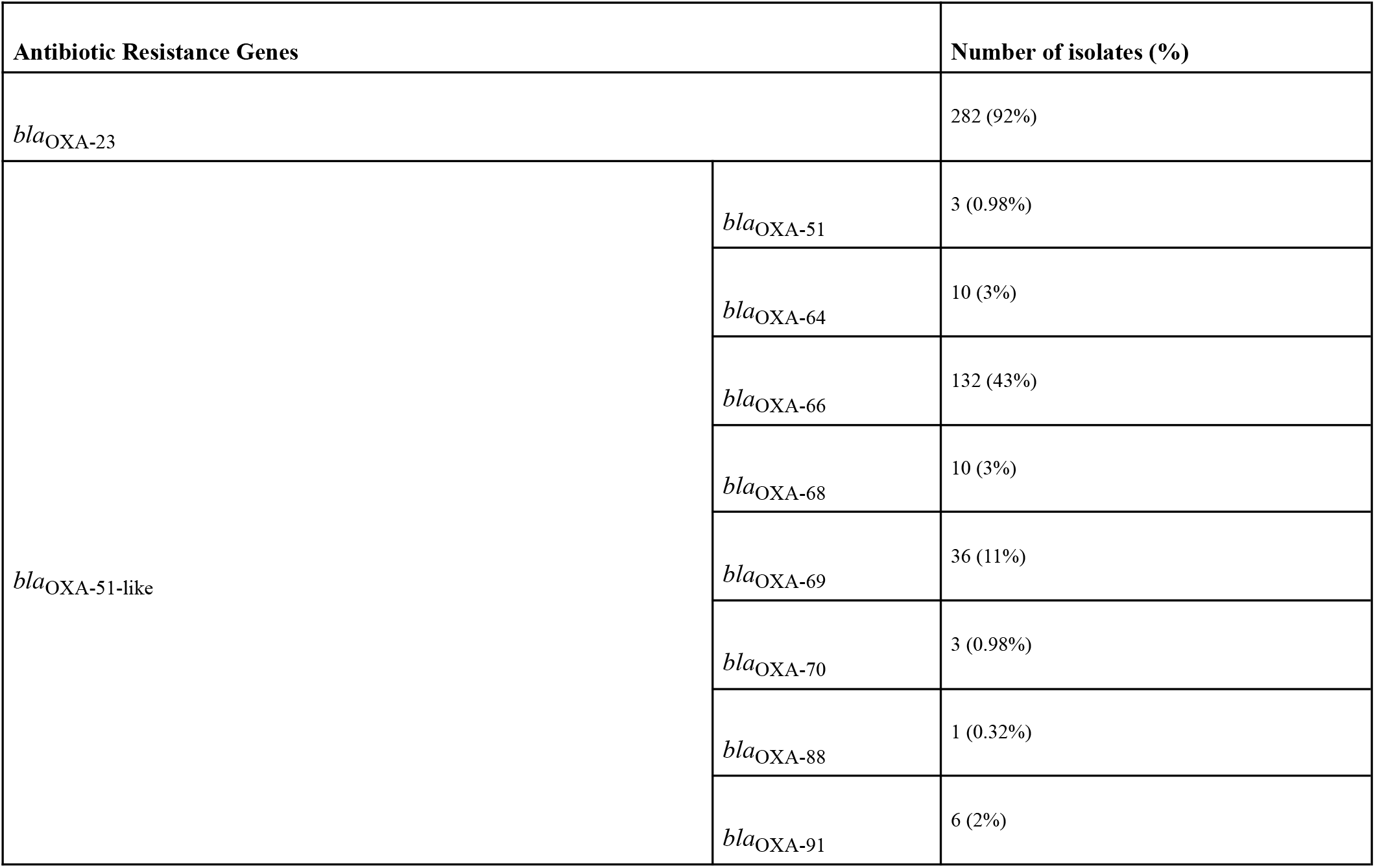
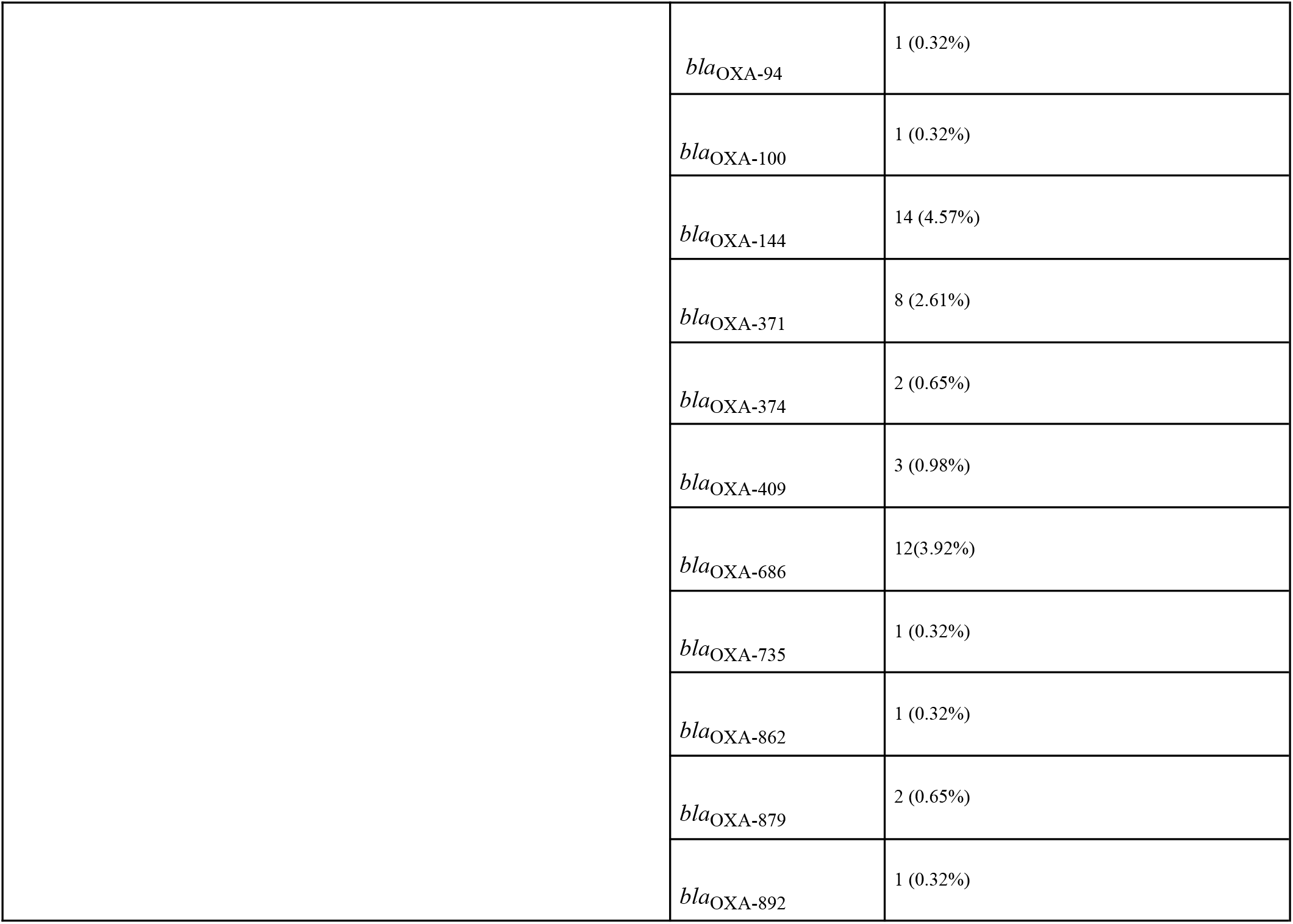
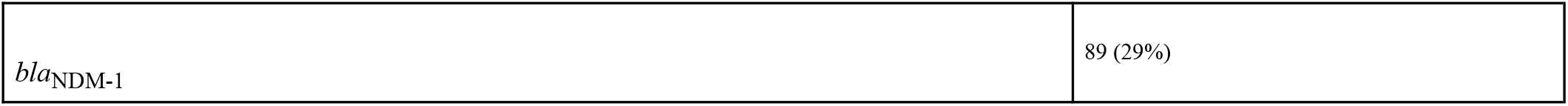
Table showing the carbapenemase encoding genes with their numbers and percentages observed in the study.

### Local outbreak of *A*.*baumannii* ST1053

WGS linked with the epidemiological data was used to construct a phylogenetic tree which clustered isolates into different clusters **(Fig 3)**. The largest cluster was observed from isolates collected from a single center, Delhi, suggesting an outbreak. All the isolates in this cluster were collected from sputum and tracheal aspirate and the patients aged between 16 - 82 years. The samples in this cluster were collected between 2018 and 2019. The MLST analysis revealed that all the isolates in this cluster belong to ST1053. Resistance profiles reported these isolates to carry *bla*_*OXA-104*_ and *bla*_*OXA-23*_ genes.

**Fig 3:**
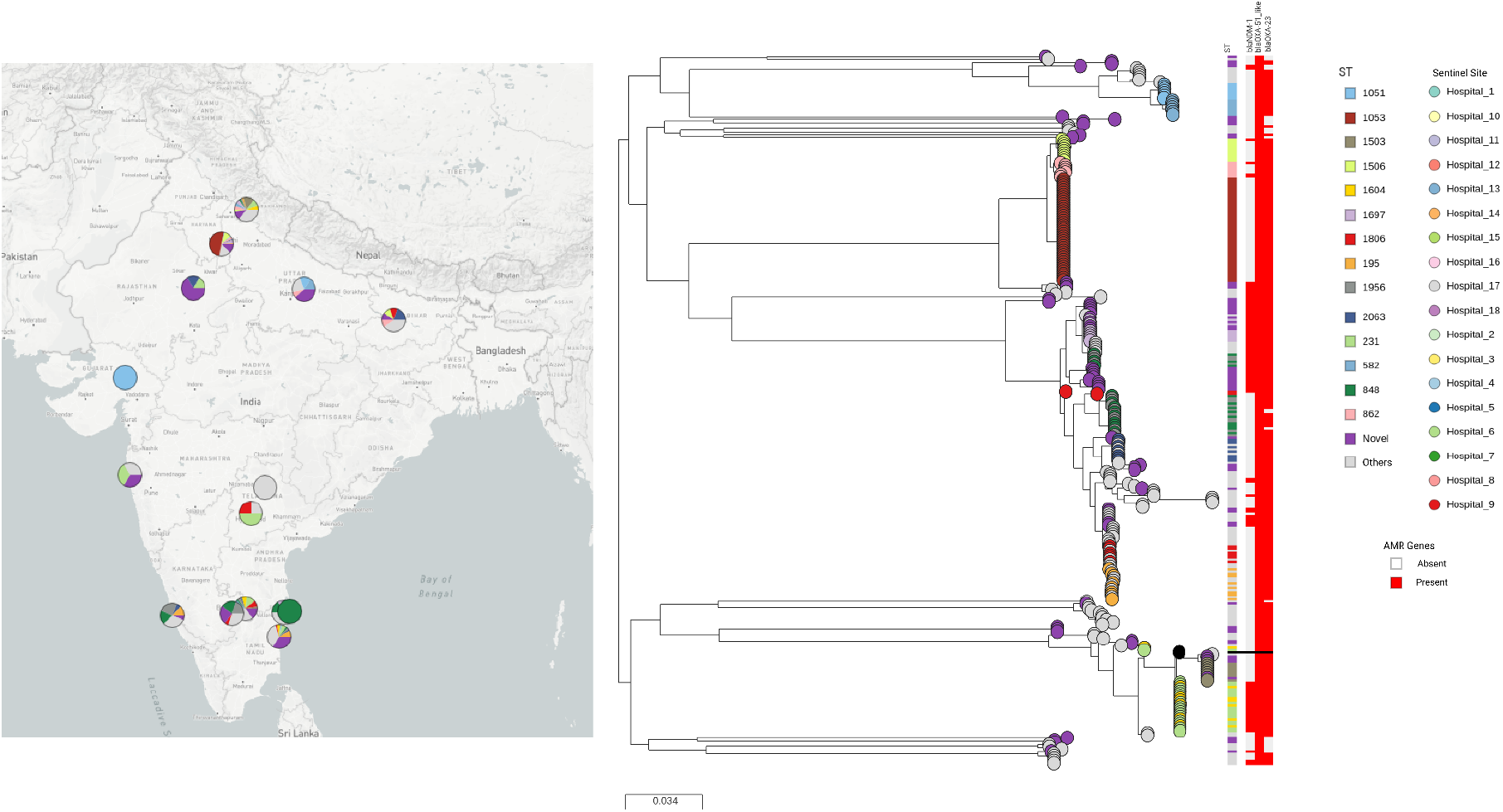
Phylogenetic tree of 306 *A*.*baumannii* Indian isolates. Tree leaves are colored by sentinel site and are annotated with ST and presence of carbapenemase genes (https://microreact.org/project/Cr9x8u93SRBbEUmQSRFLN/8f79e7ac).

## Discussion

Species of *Acinetobacter*, primarily *A. baumannii*, are becoming increasingly important nosocomial pathogens **[25]** in numerous countries across the globe including India **[26]**.The biological characteristics such as virulence, drug resistance of the clonal groups of *A. baumannii* often vary. The identification of potential reservoirs, modes of transmission is needed because of the clonal spread **[27]**. Information related to the clonal distribution across India is incomplete and there is limited published data on the STs of Indian isolates. This study offers insight into the evaluation of the molecular epidemiology of carbapenem-resistant *A*.*baumannii* in India over a period of 5 years, 2015 - 2019.

The MLST results demonstrated a significant level of diversity within the 306 *A*.*baumannii* study strains. According to the combined epidemiological and genotypic data, we identified an outbreak of *A*.*baumannii* in a single center in India based on the clusters observed in the phylogenetic tree. ST1053 was unique to this outbreak strain. Carbapenem susceptible ST1053 was first reported from Chandigarh, North India in 2014, and no reports of the same have been observed since **[28]**. The resistance profile of our study isolates revealed that they are carbapenem-resistant. It clearly shows the progression of ST1053 from carbapenem-susceptible to carbapenem-resistant from 2014 to 2019. It is also worth noting that ST1053 was not found in any other parts of India, limiting it to the country’s northern regions. We observed an absence of ST1053 reports from other countries pointing to the fact that ST1053 could be a clone specific to India. The emergence of ST1053 was investigated using goeBURST analysis on STs from the pubMLST database as well as STs from our study. The founder of ST1053 was discovered to be CC862(**Fig 4)**. SLVs and DLVs of CC862 are denoted as 1 and 2 respectively. Five STs (ST862, ST1505, ST390, ST1308, and ST1306) in this group were discovered to be specific clones in India, with no reports from elsewhere in the world.

**Fig 4:**
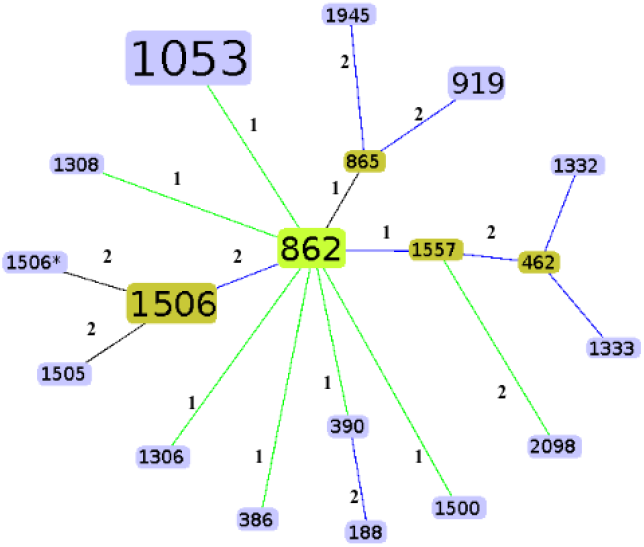
Clonal complex analysis of ST1053 isolates from the study dataset and global isolate collections from the PubMLST database.

The ST231 clone, which was shown to be dispersed among multiple specimens, was the second most prevalent clone. ST231 was initially reported internationally in Brazil in 2000, followed by reports from the United States (2005), Australia (2006), Turkey (2009), Palestine (2017) **[29]**, Egypt (2015-2017)[29], and Germany (2016). In 2014, only one study **[30]** reported the existence of ST231 in India. According to our findings, ST231 isolates were multi-centric and found in various parts of India as shown in **Supplementary Fig d**. All 17 isolates belonging to ST231 were carbapenem-resistant and carried the *bla*_NDM-1_, *bla*_OXA-23_, and *bla*_OXA-69_ genes **(Fig 2, Supplementary Table b)**.

ST1956 and ST848 were the third most common STs isolated from different specimens like blood, pus, and tracheal aspirate. ST1956 was initially discovered in the United States in 2018 and then in Sweden in 2019. These are the sole published records in the PubMLST database, and no previous publications from India were detected. We analysed the geographical distribution of ST1956 in our dataset and found it to be mostly circulating in the southern areas of India (Karnataka and Pondicherry), with a single occurrence in the North (Uttarakhand). *bla*_OXA-66_, *bla*_NDM-1_, and *bla*_OXA-23_ genes conferring resistance to carbapenems were observed in these isolates **(Fig 2, Supplementary Table b)**. ST848, which was first found in the United States in 2003, had previously been reported in another Indian study in 2015, and was the most common ST in that study **[31]**. These ST848 isolates have a similar pattern of dispersion as ST1956, with most occurrences in South India (Karnataka, Pondicherry, and Chennai) and a single incidence in the North (Uttarakhand). They were previously reported to carry *bla*_OXA-51-like_ and *bla*_OXA-23_ genes **[31]** imparting beta-lactam antibiotic resistance**[32]**. A similar observation was found in our study with ST848 isolates carrying *bla*_OXA-51-like_ and *bla*_OXA-23_ genes. Additionally, 3 isolates of each of ST1956 and ST848 also carried *bla*_NDM-1_ genes **(Fig 2, Supplementary Table b)**. Because of the similarity in epidemiological and antimicrobial resistance profiles of these two STs, we analyzed the allelic patterns of the seven housekeeping genes used for MLST typing. The only variation between the two STs was 186 nucleotide changes in a single housekeeping gene, gdhB. The allelic patterns of the remaining six housekeeping genes were all identical, suggesting that these two STs evolved together or are closely related. This was confirmed by the goeBURST analysis in which ST1956 and ST848 evolved from a common ancestor, ST451**(Fig 1)**.

The comparison of MLST results to the earlier published articles from India have been tabulated in **(Supplementary Table d.1, d.2)**. According to one study from North and South India, the most common STs were ST110, ST69, and ST103. ST848, ST451, and ST195 were the STs with the highest number of observations in another investigation **[31]**. This indicates that our analysis reported different STs to be predominant when compared to the earlier studies in India.

Molecular epidemiology of *A. baumannii* clinical isolates revealed the dominance of eight International Clonal lineages (IC-I to IC-8) all over the world. The majority of the *A. baumannii* outbreaks were linked to ICII **[33**,**34]**. In our study, Five out of 49 STs belonged to ICII (ST1806, ST208, ST195, ST218, ST848). This was in line with findings from prior Indian studies **[31]** that identified similar STs as belonging to the ICII. The emergence of ICII clones in our study represents its global expansion, which has been observed all over the world **[35]**. The most active *A. baumannii* clonal lineage is ICII, which has been linked to multidrug resistance and nosocomial infections **[36]**. The global expansion of ICII may be primarily caused by a vast clonal spread of a recent ancestor, which could be driven more internationally, implying the likelihood of long-term outbreaks. In addition, IC-I (n=3, ST1604, 231, 1503), IC-4 (n=1, ST236), and IC-8 (n=2, ST391, 1051) clones were observed in our study.

The goeBURST algorithm portrays the relationship between closely related isolates, in which the founding genotype diversifies, resulting in a cluster of closely related genotypes as its frequency in the population increases. In short, as shown by the sequence study of our Indian genotypes, there is more than one clone of *A. baumannii* circulating in India (**Fig 1)**. 32% (99/306) of the isolates belong to CC451.Among which, 37% (37/99) of the CC451 isolates were observed to be IC-II clones. Furthermore, 21% (65/306) of the isolates were clustered into CC1503, of which 10% (7/65) belong to IC-I. 20% (62/306) isolates were clustered together in CC1506 and their global clonality could not be determined. CC1604 constituted 8% (26/306) of the isolates in the study, of which 92% were found to belong to IC-I.

Carbapenem resistance among *A*.*baumannii* has been steadily increasing over the world dominated by OXA-type carbapenamase producers **[37]**. The presence of three genes, *bla*_NDM-1_, *bla*_OXA-23_, and *bla*_OXA-51-like_ genes, was found in Indian isolates, indicating a high rate of carbapenem resistance. In Indian patients, a high prevalence of the acquired *bla*_OXA-23_ *A. baumannii* strains has been observed which was in concurrence with this study where *bla*_OXA-23_ and *bla*_OXA-51-like_ genes were detected as the predominant carbapenem-resistant genes. *bla*_NDM-1_ bearing *A. baumannii* observed in our study isolates has lately been detected in a number of other countries, including Germany, Spain, Israel, Egypt, Switzerland, Libya, Pakistan, and Nepal **[38, 39, 40, 41]**. Previous research has found that the majority of isolates carrying the *bla*_NDM-1_ gene belonged to ST85, ST25, and ST222 **[42**,**43**,**44**,**45]**, isolates carrying *bla*_OXA-23_ to ST208 **[46]**, ST589 **[47]**, and isolates carrying *bla*_OXA-51-like_ genes to ST2, ST45, ST78, and ST15 **[48]**. Our study showed a difference in the pattern of STs observed in isolates harboring these genes. We found novel STs, ST1053, and ST231 as being frequent in isolates carrying these genes.

In 64 percent (197/306) of the isolates, the acquired *bla*_OXA-23_ gene coexisted with the intrinsic *bla*_OXA-51-like_ gene, while in 1 percent (4/306) of the isolates, *bla*_NDM-1_ gene coexisted with the *bla*_OXA-51-like_ gene. 27% (85/306) isolates reported co-existence of genes, *bla*_OXA-23_ and *bla*_NDM-1_ **(Fig 5)**. Co-occurrences of the *bla*_NDM-1_ gene and the *bla*_OXA-23_ gene have previously been reported in India **[49]**. Novel STs followed by ST231 and ST1604 predominated in isolates carrying *bla*_NDM-1_ and *bla*_OXA-23_ genes. 6% (20/306) of the isolates exclusively carried the *bla*_OXA-51-like_ gene **(Fig 5)**.

**Fig 5:**
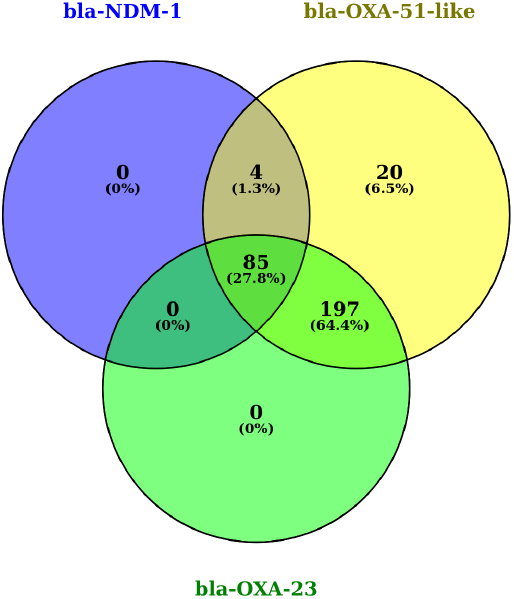
Venn diagram depicting the co-occurrences of antimicrobial genes in study isolates. The multiple existences of the genes are represented by the overlapped circle.

## Conclusion

In conclusion, WGS identified a local outbreak of ST1053 *A. baumannii* at a hospital in Northern India. India must keep an eye on this ST1053 due to its potential to cause future disease outbreaks. The antimicrobial resistance profile analysis found that *bla*_OXA*-23*_ was highly prevalent among the isolates. However, further studies are needed to identify the mechanism of acquisition of *bla*_OXA*-23*_ and *bla*_NDM-1_ genes in our isolates.

## Acknowledgements

Members of the NIHR Global Health Research Unit for the Genomic Surveillance of Antimicrobial Resistance

